# Highly multiplexed, image-based pooled screens in primary cells and tissues with PerturbView

**DOI:** 10.1101/2023.12.26.573143

**Authors:** Takamasa Kudo, Ana M. Meireles, Reuben Moncada, Yushu Chen, Ping Wu, Joshua Gould, Xiaoyu Hu, Opher Kornfeld, Rajiv Jesudason, Conrad Foo, Burkhard Höckendorf, Hector Corrada Bravo, Jason P. Town, Runmin Wei, Antonio Rios, Vineethkrishna Chandrasekar, Melanie Heinlein, Shuangyi Cai, Cherry Sakura Lu, Cemre Celen, Noelyn Kljavin, Jian Jiang, Jose Sergio Hleap, Nobuhiko Kayagaki, Felipe de Sousa e Melo, Lisa McGinnis, Bo Li, Avtar Singh, Levi Garraway, Orit Rozenblatt-Rosen, Aviv Regev, Eric Lubeck

## Abstract

Optical pooled screening (OPS) is a highly scalable method for linking image-based phenotypes with cellular perturbations. However, it has thus far been restricted to relatively low-plex phenotypic readouts in cancer cell lines in culture, due to limitations associated with *in situ* sequencing (ISS) of perturbation barcodes. Here, we developed PerturbView, an OPS technology that leverages *in vitro* transcription (IVT) to amplify barcodes prior to ISS, enabling screens with highly multiplexed phenotypic readouts across diverse systems, including primary cells and tissues. We demonstrate PerturbView in iPSC-derived neurons, primary immune cells, and tumor tissue sections from animal models. In a screen of immune signaling pathways in primary bone marrow-derived macrophages, PerturbView uncovered both known and novel regulators of NFκB signaling. Furthermore, we combined PerturbView with spatial transcriptomics in tissue sections from a mouse xenograft model, paving the way to *in vivo* screens with rich optical and transcriptomic phenotypes. PerturbView broadens the scope of OPS to a wide range of models and applications.

## INTRODUCTION

Pooled screens have had a transformative effect on our ability to link genetic perturbations with their phenotypic outcomes at scale. While historically limited to a bulk readout of a few parameters^1,2^, typically through cell enrichment by viability or FACS, pooled screens have recently been integrated with high-dimensional phenotypic readouts through molecular profiling^3–5^ (Perturb-Seq) or optical imaging^6^. These approaches have made it possible to study gene function and regulatory circuits at unprecedented resolution. In particular, imaging by microscopy is a highly versatile screening tool with a wide range of phenotypic assays that can be tailored to specific biological questions. Several recent approaches have enabled image-based screening with pooled libraries, relying on 1) selection of cells-of-interest by imaging, followed by bulk sequencing of perturbation barcodes^7–11^, or 2) optical measurements of phenotypic markers along with nucleic acid- or protein-based perturbation barcodes in each cell^12–15^. In this latter class, optical pooled screens (OPS) use targeted *in situ* sequencing (ISS) of perturbation barcodes to perform highly scalable imaging screens. OPS has been used to study perturbation effects on immune signaling^13^, viral infection^16^, the unfolded protein response^17^, and cell morphology^18–21^.

However, OPS has thus far been limited to relatively low-plex molecular measurements in cancer cell lines, largely due to challenges in robustly detecting perturbation barcodes by ISS. Non-cancer cell types, especially certain primary cells, notoriously express barcodes at low levels^22^, making it difficult to assign perturbations to cells. In addition, many highly multiplexed imaging technologies rely on iterative rounds of staining and imaging^23,24^ that are incompatible with the inherent fragility of RNA barcodes, which are susceptible to washing or degradation.

Here, we address these limitations by developing PerturbView (<u>Perturb</u>ation experiments with a <u>V</u>isual *<u>I</u>n vitro* transcription <u>E</u>nabled <u>W</u>orkflow), an approach that leverages *in vitro* transcription (IVT) for robust *in situ* sequencing of perturbations across diverse cell types and tissues. PerturbView expands on the Zombie protocol^25^, which uses phage RNA polymerase (RNAP) to express RNA barcodes after cell fixation, decoupling perturbation detection from barcode expression, and simplifying the integration with iterative molecular phenotyping. We designed and validated a PerturbView vector that is compatible with Perturb-Seq^3^, so that the same cell libraries can be profiled by both approaches; demonstrated robust barcode detection across multiple cell models, including primary cells and iPSC-derived cells; and conducted a focused pooled CRISPR screen for regulators of p65 translocation in primary bone marrow-derived macrophages (BMDMs), recovering both established and novel hits. PerturbView also enabled a multimodal screen in primary macrophages by combining hybridization chain reaction fluorescent *in situ* hybridization (HCR FISH)^26^ and immunofluorescence by IBEX^27^ with ISS of perturbations. Finally, we integrated PerturbView readout of barcodes expressed *in vivo* in a cancer xenograft model with spatial transcriptomics to perform rich expression profiling in a native tissue environment. PerturbView significantly broadens the utility of OPS by making it possible to screen across diverse biological contexts with rich phenotypic readouts, enhancing our ability to decipher the functions of genes and regulatory elements in both intracellular and intercellular circuits.

## RESULTS

### PerturbView enables robust and rapid barcode detection for pooled optical CRISPR screens

Because many primary cells express barcodes at relatively low levels (**Fig. S1a**), we developed PerturbView, which enables facile optical detection of sgRNAs by utilizing T7 RNAP to amplify chromosomally integrated CRISPR perturbation vectors after cell fixation and phenotyping (**Fig. 1a**). We hypothesized that the versatility and robustness of OPS could be improved by leveraging the recently developed Zombie protocol^25^ to decouple barcode expression from barcode detection. The large molecular amplification of T7 RNAP produces a massive number of sgRNA molecules in the nucleus, which can be easily detected by *in situ* sequencing. This addresses the limitations of the conventional OPS protocol in detecting perturbations from cells with low levels of RNA barcode expression or barcode loss during phenotyping.

**Fig. 1.**
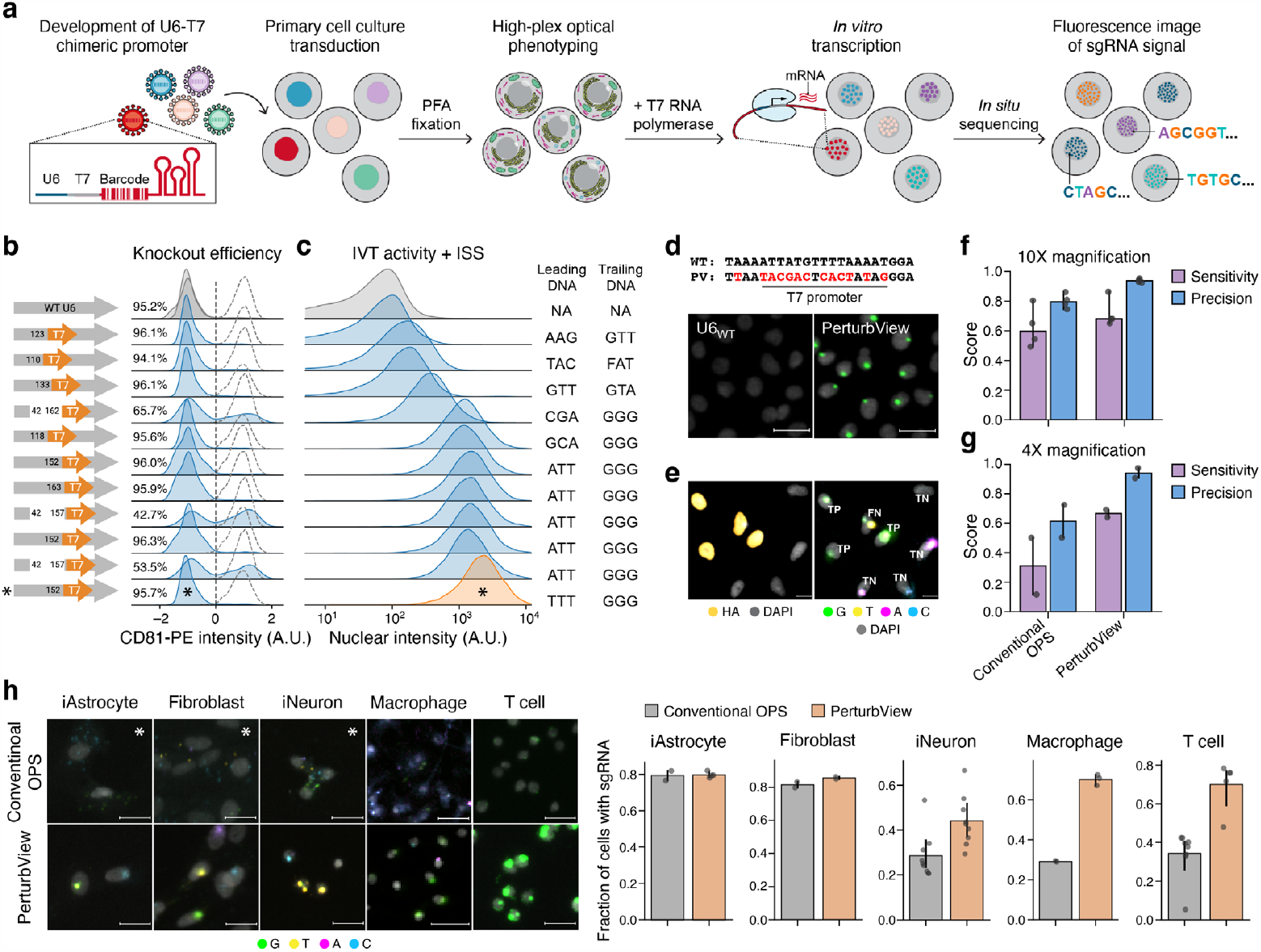
PerturbView. (**a**) Method overview. (**b**,**c**) Joint optimization of CRISPR efficiency and IVT activity. Distribution of CD81 expression levels (b; control-normalized *Z*-scores) and IVT+ISS nuclear intensity (c; single cell fluorescence intensity of sgRNA reads from each promoter variant) for each promoter design (b, left; numbers indicate the position of the T7 promoter insertion and U6 truncations with respect to the U6 promoter start) in A549 cells. Top: a standard CROPseq CD81 sgRNA (gray). Dotted lines: CD81 distribution in wild-type (WT) cells. Knockout efficiency is labeled as % population below 0 (vertical line). At least *k* = 5,517 cells per promoter; 10,236 on average. Asterisks: chosen PerturbView construct. (**d**) PerturbView enables bright IVT from the U6 promoter with minimal modifications to the U6 sequence. Top, WT U6 promoter and PerturbView construct sequences. Bottom, A549 cells post IVT from the original CROP-Seq and PerturbView vector (Green = G base, scale bar, 50 μm). (**e**) Frame-shift reporter screen to determine sensitivity and specificity of barcode detection. Left, Representative image of cells with IF of HA-staining (right) or guide decoding with screen assignments (left, TP, FN and TN labeled by presence or absence of HA epitope and the first base of reads) (scale bar, 25 μm). (**f**,**g**) Improved sensitivity and precision of PerturbView. Sensitivity (purple) and precision (blue) of barcode detection (y axis) by conventional ISS-based sgRNA detection and PerturbView (x axis) at 10X (f) and 4X (g) magnification (*n* = 4 independent replicates; SD error bar, each dot representing a replicate). (**h**) PerturbView performs well across primary and iPSC-derived cells. Left, Representative images of sgRNA barcode detection by conventional ISS and PerturbView across cell types (Scale bar, 25 μm). All images are acquired identically except images denoted by an asterisk (* denotes a 4-fold longer exposure duration to account for the drastically dimmer signal of some cell types in standard ISS experiments). Right, Fraction of cells positive for sgRNA signal (>1 sgRNA read per cell, mean of well replicates with error bar indicating SD, each dot representing a replicate) across cell types. *n* ≥ 2 for iAstrocytes (mean *k* = 2,107 cells per replicate), *n* = 2 for IMR90 fibroblasts (*k* = 67,547 cells), *n* = 9 for iNeurons (*k* = 15,332 cells), *n* ≥ 2 for immortalized macrophages (*k* = 175,182 cells), *n* ≥ 5 for T cells (*k* = 4518 cells).

We first showed that barcodes can be robustly amplified and read through IVT and ISS in fixed cells. To this end, we tested three vectors: the CROP-seq^28^ vector, and two CROP-seq derivatives, where the T7 promoter was placed either upstream of or in place of the U6 promoter (**Fig. S1b**). We transduced the constructs into A549 cells, performed antibiotic selection, seeded cells onto glass-bottom plates, and fixed the sample with paraformaldehyde (PFA), a common fixative that is compatible with many image-based phenotypic readouts. To avoid inhibition of IVT by PFA fixation^25^, we added a decrosslinking step^29^, followed by sgRNA overnight amplification with T7 RNAP (Methods). Following IVT, we performed the conventional OPS protocol^21,30^ and sequenced the first base of the sgRNA. Constructs containing the T7 promoter produced sgRNA signal as bright nuclear foci (**Fig. S1b**). Amplification was lower when the T7 promoter was placed upstream of the U6 promoter, leading us to hypothesize that the distance between a T7 promoter and a barcode is inversely related to IVT efficiency, likely due to reduced polymerase access to DNA in the presence of cross-links.

In order to achieve both high CRISPR and IVT efficiencies, we designed a chimeric promoter, where the T7 sequence is embedded within the U6 promoter as close as possible to the sgRNA without disrupting CRISPR activity. The published U6/ T7 promoter design^31,32^ resulted in substantial decreases in editing efficiency (**Fig. S1c,d**), necessitating the design of a new chimeric promoter to support highly efficient screens. We tested 13 vector designs, varying the location of the T7 promoter sequence as well as the leading/trailing nucleotides^33^ (Methods, **Table S1**). To assess which chimeric promoter maintains high CRISPR editing activity, we evaluated CRISPR editing efficiency using a CD81 flow cytometry assay in A549 cells (Methods), and identified several chimeric promoters with comparable CRISPR performance to the CROP-seq vector (**Fig. 1b**). We note that truncated U6 variants based on the mini U6 promoter^34^ resulted in decreased editing activity (**Fig. 1b**).

Among the chimeric promoters, we selected an optimal construct that supported both efficient CRISPR activity and successful sgRNA detection by OPS after IVT (**Fig. 1b,c, asterisk**). The chimeric promoters varied >40-fold in nuclear sgRNA production by IVT (**Fig. 1c**; 62±66 to 2,608±1,898 AU; min to max mean nuclear signal ± standard deviation (SD)), with AT-rich leading sequences and trailing G triplet crucial for optimal T7 activity, as reported previously^33^ (**Fig. 1c**). The chosen construct (**Fig. 1b, asterisk**), used for all further PerturbView screens, had a 13 bp replacement from the original U6 promoter and high IVT efficiency (**Fig. 1c,d**), while retaining CRISPR editing efficiency in all tested cell types (**Fig. 1b, Fig. S1e**).

We observed a tradeoff between the sensitivity and precision of perturbation detection that we tuned by varying decrosslinking and IVT parameters. In particular, higher levels of decrosslinking and IVT can increase signal levels, but could also result in false positives due to the diffusion of barcode RNAs into neighboring cells^30^. In order to optimize this tradeoff, we performed a miniature screen with a frameshift reporter assay^13^ to quantify both genotyping efficiency (barcode detection, i.e., ‘sensitivity ‘) and phenotype-to-genotype mapping precision. We cloned a pool of targeting and non-targeting sgRNAs into the PerturbView vector, where each targeting sgRNA can lead to a frameshift and the expression of an HA-epitope fused to histone H2B. In order to calculate sensitivity and precision of barcode detection, we categorized each cell in the experiment as follows: HA-positive cells with a detected targeting sgRNA are true positives; HA-positive cells with a non-targeting sgRNA or without a detected sgRNA are false negatives, HA-negative cells with targeting sgRNA are false positives, and HA-negative cells with non-targeting sgRNA or without a detected sgRNA are true negatives (**Fig. 1e**). Using this system, increasing heat decrosslinking from 0–1 hour to 4 hours increased both sensitivity and precision, with no further improvement beyond 4 hours (**Fig. S1f**), confirming the importance of decrosslinking. Next, even short IVTs of 3 hours had comparable sensitivity and precision as longer (24 hr) durations (**Fig. S1g**). Thus, we chose 4-hour decrosslinking and 4-hour IVT for PerturbView, achieving improved sensitivity (72±9.7%; mean±SD) and precision (94±1.3%) (in A549 cells) compared to conventional OPS (62±3.5% sensitivity and 80±15.6% precision) (**Fig. 1f**).

The high levels of signal amplification in PerturbView compared to conventional OPS make it possible to boost data capture rates, which are currently a major bottleneck in large screens. While standard OPS typically requires imaging at 10X or higher magnifications to capture the ISS signal, PerturbView yields mappable reads at 4X magnification. Performing the frameshift reporter assay at 4X magnification achieved comparable results to 10X magnification (67% sensitivity and 94% precision) (**Fig. 1g, Fig. S1h**). For a typical genome-wide CRISPR screen, which usually requires 4–10 plates, this reduces the imaging time for 12 cycles of sequencing from 24 hours (10X) to ∼ 4 hours (4X) per plate.

### PerturbView is compatible with primary and iPSC-derived cells

PerturbView enabled screening in diverse cell types, in contrast to conventional OPS, which can be challenging to perform in non-cancer cell lines due to poor sgRNA detection. To demonstrate this, we tested PerturbView across iPSC-derived neurons (iNeurons), iPSC-dervied astrocytes (iAstrocytes), primary human fibroblasts (IMR90), primary human T cells and immortalized mouse macrophages (Methods). In fibroblasts and astrocytes, where conventional OPS was relatively efficient, PerturbView showed comparable performance (**Fig. 1h**). Conversely, in iNeurons, macrophages and primary T cells, where conventional OPS performs poorly, PerturbView substantially improved barcode detection (1.6-, 2.9- and 2.0-fold, respectively, **Fig. 1g**).

### Dissection of NFκB signaling in primary macrophages with PerturbView

NFκB signaling is a critical pathway that cells use to respond to stresses and stimuli. While OPS of the pathway has been performed in a cancer cell line, it is especially crucial in primary immune cells, and thus screening it in this native context may highlight distinct functions. To this end, we conducted a PerturbView screen for regulators of p65 translocation^13^ in primary mouse bone marrow-derived macrophages (BMDMs) (**Fig**.**2a**). We constructed a PerturbView CRISPR sgRNA library, targeting 163 genes with 4 sgRNAs per gene, including 10 non-essential control genes, along with 60 non-targeting control sgRNAs (Methods). The library is composed of components of the canonical NFκB pathway, along with various other metabolic and signal transduction genes (**Table S1**). Seven days after sgRNA transduction, we stimulated the transduced BMDMs with either TNFα, IL-1β, LPS or a H_2_O control for 40 minutes, fixed and stained for p65, and performed ISS by the conventional method (without T7 IVT) and by PerturbView (with T7 IVT). Conventional OPS efficiency was dramatically reduced by p65 phenotyping, even when using RNase inhibitors during staining and imaging (from 39% to 11%; **Fig. 2b**), while PerturbView after p65 immunofluorescence showed a 80% barcode detection rate (**Fig. 2b**). sgRNA representation by PerturbView was highly concordant with that from next-generation sequencing of genomic DNA extracted from the cell library (**Fig. S2a**). Moreover, PerturbView also offered greater flexibility than conventional OPS, because ISS needs to be processed immediately after fixation^30^, whereas genotyping efficiencies with PerturbView were similar between fresh plates immediately post-fixation, and experiments performed one week following fixation (**Fig. 2b**, grey vs. red circles). We thus focused on PerturbView results for all further NFκB screens, scoring each gene (4 guides per gene) based on nuclear p65 levels (median of all guides per gene) (**Fig. 2c–e, Table S1**, Methods).

**Fig. 2.**
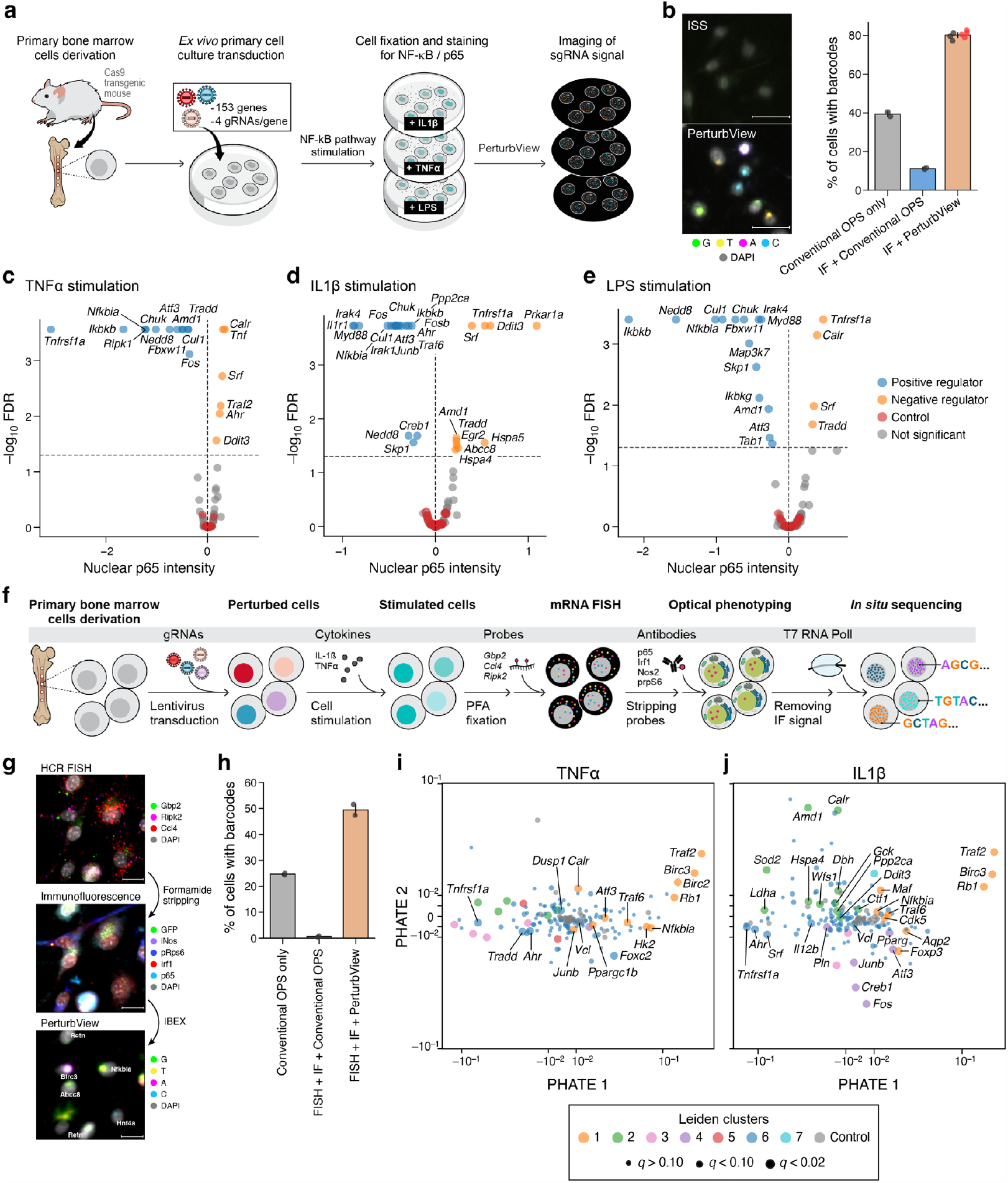
PerturbView screens with multiplex molecular readouts in primary immune cells reveal shared and context-specific mechanisms of NFKB signaling. (**a**) Overview of NFKB screen in primary mouse BMDMs. (**b**) PerturbView but not conventional OPS sensitively detects sgRNA barcodes in BMDMs. Left, Reads (fluorescent signals) from standard ISS (top) and PerturbView (bottom) in BMDMs (Scale bar, 25 μm). Right, Mean fraction of cells with detectable reads (y axis) from standard ISS (conventional OPS, *n* = 2 technical replicates), standard ISS after IF (*n* = 2 technical replicates) and PerturbView after IF (*n* = 2 replicates pooled from three independent experiments. Red dots: Replicates phenotyped and genotyped a week after fixation). Error bars: SD. (**c–e**) Shared and context-specific hits from PerturbView OPS in three stimuli. Significance (-log10(FDR), y axis) and effect size (normalized nuclear p65 intensity) for each perturbation (dot) in TNFα (c), IL-1β (d), and LPS (e) stimuli, colored for positive (blue; FDR < 0.05), negative (orange; FDR < 0.05), control (red, non-targeting sgRNA and non-essential genes) perturbations. (*k* = 1940±541, 2874±809, and 990±288 (mean±SD) cells per targeting gene in TNFα (c), IL-1β (d), and LPS (e). (**f**) Workflow for multimodal NFKB PerturbView screen using RNA FISH (HCR) and IF (IBEX). (**g**) Representative image of BMDM assayed in order by HCR FISH (top), immunofluorescence (middle), and PerturbView (bottom). (**h**) Successful sgRNA barcode detection in multimodal PerturbView, but not conventional OPS. Mean percent of cells with detected barcode (*y* axis) in conventional OPS, conventional OPS after FISH and IF, and multi-modal PerturbView (*x* axis). Error bars: SD. Dots: each well and condition. (**i,j**) Co-functional perturbation modules based on multimodal PerturbView. PHATE embedding of perturbation profiles (dots) following TNFα (i) or IL-1β (j) stimulus, sized by q value of perturbation effect and colored by cluster or as controls. Gene labels are shown for *q*-value < 0.02 in selected clusters (1 and 6 in (i); 1, 2, 4 and 6 in (j)).

Our screen recovered established NFκB regulators as hits, including Ikbkb, Nfkbia, Tnfrsf1a, Chuk, Nedd8, Tradd, Irak4, Myd88, Ripk1, Traf2, Traf6, Irak1, and Il1r1, in the correct, stimulus-dependent manner, and captured both positive and negative regulators (which reduce or increase p65 signals when perturbed, respectively) (**Fig. 2c–e**). While some NFκB regulators played the same role in all three conditions, many others were context-dependent, playing a role only under relevant stimuli (**Fig. 2c–e** and **Fig. S2b**). For example, Tnfrsf1a was the strongest hit for TNFα stimulation and Il1r1 for IL1β stimulation35; Myd88 and Irak4 mediated both IL1β and LPS signaling but not TNFα signaling^35^.

Notably, Map3k7 (Tak1), a core NFκB component, only impacted p65 nuclear levels under LPS stimulation, but not under TNFα or IL1β stimuli (**Fig. S2c**), suggesting that Map3k7 is only required for NFκB activation in some contexts^36,37^. Prior screens conducted in human cancer cell lines^13^ showed that Map3k7 is necessary for both TNFα and IL1β responses, whereas a screen in primary mouse bone-marrow-derived dendritic cells (BMDCs) showed that Map3k7 is dispensable for TNF protein production following LPS signaling in DCs^38^. This highlights the context specificity even of key components of the NFκB pathway, in terms of both stimulus and cell type context, and thus the importance of PerturbView ‘s ability to screen in diverse primary cells.

Although our screen is primarily designed for loss-of-function phenotypes (leading to a reduction in p65), PerturbView also identified several negative regulators (resulting in a p65 increase), a more challenging task^38^, given the already high p65 signals in the unperturbed, stimulated cells. These included Prkar1a, where KO increased the response to IL1β, and Tnfrsf1a KO, which led to the expected loss of p65 signal under TNFα stimulus, but increased the p65 response in both LPS and IL1β (**Fig. S2d,e**). Thus, PerturbView provided a generalizable, robust approach for image-based pooled screening across diverse, biologically relevant cell contexts.

### Joint RNA and protein profiling uncovered NFκB regulatory modules

Multiplexed imaging, including iterative immunofluorescence^27,39–41^ and highly multiplexed FISH^42–44^, can provide systematic discovery tools for OPS screens, analogous to single-cell RNA-seq (scRNA-seq) profiles^3^, but with the added richness of cell biological imaging phenotypes. However, thus far, OPS has been limited to phenotyping with a few markers (up to 7 shown to date^16^), requiring specialized staining and bleaching procedures^19,20^ and careful interleaving of the phenotypic assay with the ISS workflow. This is primarily because phenotyping assays often interfere with ISS efficiency and vice versa^30^, resulting in a tradeoff between phenotypic data quality and perturbation detection (as shown above for even simple immunofluorescence with conventional ISS, **Fig. 2b**). This is especially challenging for assays that require multiple cycles of staining and imaging, with each cycle increasing the probability of RNA loss.

PerturbView addresses this challenge by making it possible to perform all phenotyping steps upstream, prior to barcode RNA production from IVT and ISS. We leveraged this capability to enhance the information content of our NFκB stimulation screens in BMDM with joint RNA and protein profiling (**Fig. 2f**). Specifically, after 24 hours of IL1β or TNFα stimulation of BMDMs (perturbed with the same library as before), we fixed the cells and performed HCR FISH to measure three key transcripts (Gbp2, Ccl4 and Ripk2) followed by immunofluorescence for four proteins (Irf1, Nos2, p65 and phospho-rpS6). These genes are regulated by inflammatory stimuli, and were chosen to cover diverse functions, such as intracellular (Irf1, p65) signaling, intercellular (Ccl4) signaling and immunometabolism (Gbp2, Nos2 and phospho-rpS6). We stripped the FISH probes with formamide treatment and bleached immunofluorescence signals with lithium borohydride prior to IVT and ISS (Methods, **Fig. 2g**)^27^. These approaches are compatible with high-plex profiling^45^ via further steps of iterative staining and signal quenching, paving the way to PerturbView screens with highly multiplexed molecular profiling readouts. While PerturbView successfully recovered phenotypes and perturbations in 49.5% of BMDMs, conventional OPS was only successful in 0.6% of cells after phenotyping, underscoring the challenges of integrating complex assays with conventional ISS (**Fig. 2h**).

Combining perturbations with high-dimensional, multiplexed spatial features in this PerturbView screen enabled a richer dissection of perturbation effects, such as deciphering co-functional modules of perturbed genes based on their shared phenotypic profiles. To this end, for each guide, we extracted features associated with RNA, protein localization and cell morphology. We filtered and reduced the features to 50 principal components, and aggregated them for each guide by taking the median across cells (Methods). We embedded the aggregated feature profiles into a PHATE space^46^. First, many guide replicates targeting the same gene grouped together (**Fig. S2f,g**) in the relevant conditions (e.g., all four guides targeting Tnfrsf1a in the TNFα but not IL1β stimulus; **Fig. S2f**). Consistently, different guides targeting the same gene had higher cosine similarity of their median phenotypic profiles than guides targeting random pairs of genes, whereas guide pairs targeting genes unexpressed in BMDMs (Olfr genes) did not, suggesting impactful and consistent phenotypic effects for the targeting guides (**Fig. S2g**).

Next, embedding the gene-level profiles under TNFα and IL1β stimulation into a single PHATE space (**Fig. 2i,j, Table S1**), revealed perturbation profiles that grouped in ways reflecting both shared and unique features of TNFα and IL1β signaling. For example, while profiles for Tnfrsf1a and Tradd (**Fig. 2i**, cluster 6), the protein complex that initiates TNFα signaling, were closely grouped in the TNFα stimulation screen, this co-functional relation was not observed under IL1β stimulation (**Fig. 2j**). Similarly, Birc2 (cIAP1), Birc3 (cIAP2) and Traf2, which also form a protein complex mediating TNFα signaling, group together in TNFα stimulation (**Fig. 2i**, cluster 1), but in IL1β, only Traf2 and Birc3 group together, along with other NFκB components, Nfkbia and Traf6. Neither Birc2 nor Birc3 impacted p65 translocation (**Fig. 2c,d**), suggesting that they mediate downstream immune signaling. A module observed only in IL1β signaling (**Fig. 2j**, cluster 4) includes Atf3, Fos, Creb1, Junb and Pln: Atf/Creb family factors are known to interact with AP-1 proteins, such as Fos and Junb, and to regulate Pln47. A key metabolic module was also unique to IL1β stimulation, including Ldha, Hspa4, Ddit3, Calr and Gck, which mediate mTOR and hypoxic signaling^48^ and may reflect HIF1α-mediated metabolic reprogramming during macrophage activation^49^ after IL1β stimulation.

Finally, we assessed how each transcript and protein contributes to a cell ‘s phenotype. To this end, we restricted the input features to only one of eight (each of the seven molecular species and DAPI), scored each of them for impact (Methods), and clustered perturbations by their score profiles across individual features. This allowed us to identify which phenotypic features yield similar perturbation scores, and which features contribute to the ‘impact’ call of each perturbation and in which condition (**Fig. S2h**). For example, in TNFα stimulation, phospho-rpS6 was associated with the impact of perturbations in Maf, Traf2, Ldha and Sod2, and quite distinct from all other phenotypic features. In IL1β stimulation, Ccl4 contributed to calling a separate set of impactful perturbations than Ripk2, Gbp2, or other features, consistent with a recent Perturb-seq study^50^ reporting Ripk2 and Gbp2 as part of the same gene program, distinct from Ccl4. Taken together, PerturbView allows highly multiplexed imaging screens with RNA and protein multiplex readouts in primary immune cells.

### PerturbView enables optical tissue perturbation screens along with spatial transcriptomic readout

Directly performing pooled perturbation screens in tissue holds vast potential for understanding tissue biology, while reducing the number of animals required for characterizing complex organismal biology. However, *in situ* spatial tissue perturbation experiments are technically challenging, because the difficulty of detecting a sgRNA is further compounded by the inefficiency of performing assays in tissue.

To demonstrate how these limitations are overcome by the amplification of sgRNA barcodes through PerturbView, we delivered a pool of 100 barcoded (non-targeting) sgRNAs to DLD-1 colon cancer cells, transplanted the cells into mice to generate a xenograft tumor, and collected the tumor to perform IVT and ISS in fresh-frozen and formalin-fixed paraffin-embedded (FFPE) sections. With PerturbView, we observed bright, discernible foci, which unambiguously corresponded to sgRNA after ISS (**Fig. 3a**). We successfully sequenced barcodes over six rounds of decoding, yielding a >90% assignment rate of ISS reads to sgRNAs (**Fig. S3a**). Of 100 barcodes transduced to DLD-1 cells, 66 and 90 were retrieved by PerturbView (with at least 10 reads per experiment replicate) in the FFPE and fresh frozen samples, respectively (**Fig. 3b**). The fraction of cells with identified barcodes (with ≤1 Hamming distance) was 40% in fresh–frozen sections and 16% in FFPE sections (**Fig. 3c**, note that only some of the cells in the tumor are barcoded human cells; the rest are mouse cells). sgRNAs representation agreed well with that from next-generation sequencing of the plasmid library (**Fig. 3d**; Pearson ‘s *r* = 0.79, 0.76 for FFPE and fresh–frozen tissue, respectively). Thus, PerturbView is viable for performing guide detection *in situ* for tissue screens.

**Fig. 3.**
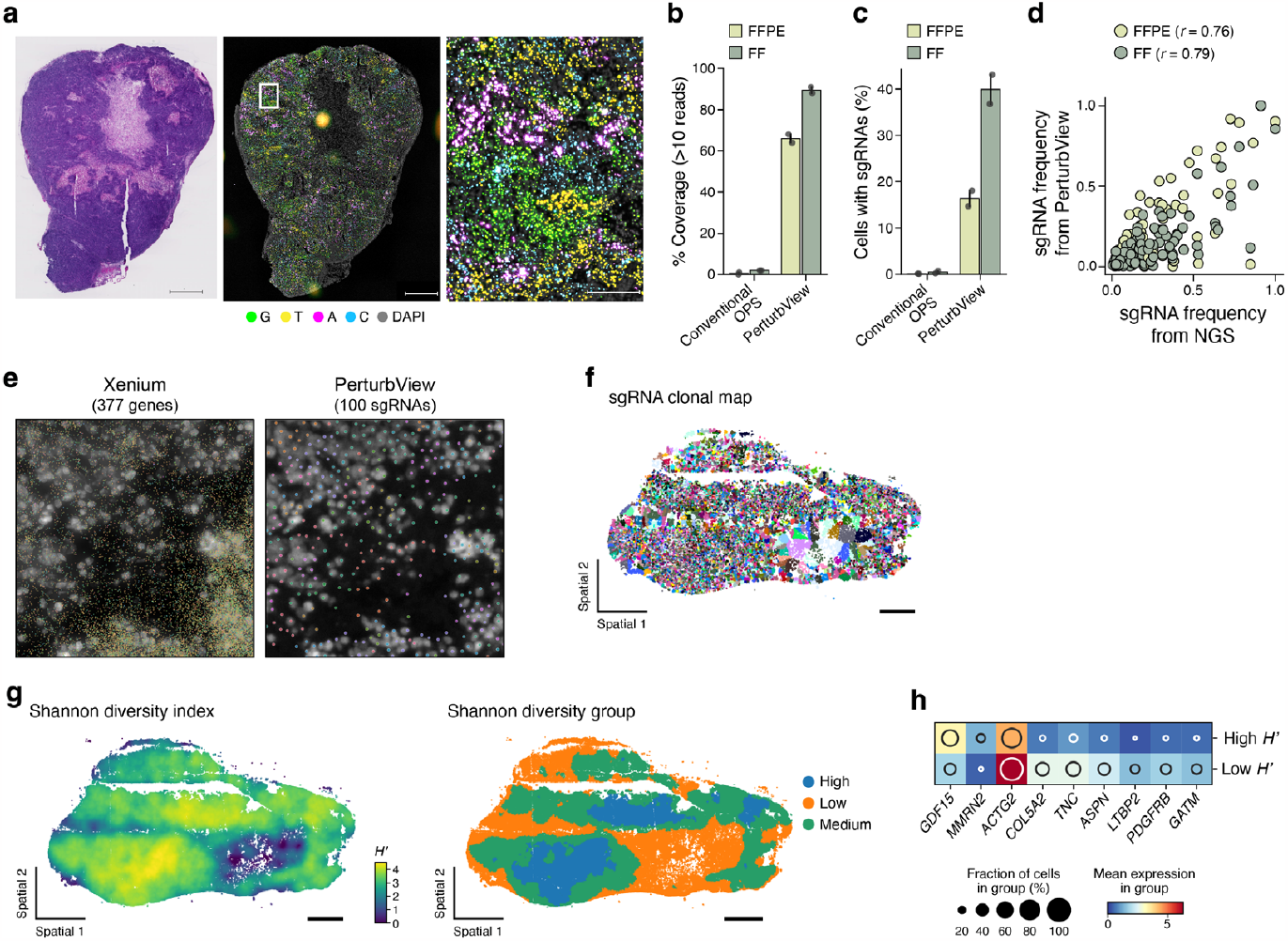
PerturbView for sgRNA detection and spatial transcriptomics in tissue. **(a–d)** PerturbView facilities pooled sgRNA detection in tissue. (a) Representative H&E staining image (left, scale bar = 1mm) and PerturbView detection of sgRNAs (right, colored bases scale bar = 1 mm) of a DLD-1 subcutaneous xenograft. Right, single cell PerturbView detection in an inset zoom (scale bar = 200 μm). (b,c) Mean percent of sgRNA library (b, y axis) and mean percentage of cells with detectable sgRNAs (c, y axis), captured by standard ISS or by PerturbView (x axis) in FFPE (light green) or fresh frozen (FF) (dark green) subcutaneous tumors. *n* = 2 independent experiments, Error bars: SD. (d) Frequency of each sgRNA (dots; normalized counts) estimated by NGS of the plasmid library (*x* axis) or PerturbView (*y* axis) of FF (dark green) or FFPE (light green) samples. *n* = 2 for each FF and FFPE treatment. Pearson ‘s r is noted. (**e–h**) PerturbView combined with spatial transcriptomics relates gene expression to clonal identity. (e) Tissue section image of Xenium spatial transcriptomics (left; 377 human pan-cancer markers) followed by sgRNA identification with PerturbView (right). (f) Tumor section colored by sgRNA clone (scale bar, 1 mm). (g) Shannon index computed from clonal analysis (g, left) or thresholded into low, middle and high Shannon diversity (g, right) (scale bar, 1 mm). (h) Genes (columns) that are differentially expressed between regions with high and low Shannon diversity (rows, shared hits between *n* = 2 tumors with 2 tissues sections per tumor).

Next, we combined PerturbView with spatial transcriptomics, leveraging Xenium technology^51^ for fresh–frozen tissue with a multi-tissue and cancer gene panel (Methods). Following the Xenium assay and DAPI staining, we performed PerturbView and captured sgRNA sequences over six cycles of ISS (**Fig. 3e**, Methods). PerturbView successfully produced joint RNA– sgRNA maps. After the Xenium run, ∼100% of ISS reads were mapped with high fidelity (within ≤2 Hamming distance) to sgRNA barcodes (**Fig. S3b**).

Finally, we leveraged our spatial PerturbView data sets to ask what tumor intrinsic gene programs were associated with clonality in our tumor model, hypothesizing that specific cellular expression patterns were associated with the growth, survival and migration of tumor cells. Using the non-targeting sgRNAs as high-diversity clonal barcodes (descendents of transplanted cells all expressed the same sgRNA), we generated clonal maps of each sgRNA throughout the tumors (**Fig. 3f**). We then assigned tissue regions into three classes, based on the local clonal diversity of barcodes by Shannon ‘s diversity index H ‘: low (bottom 25th percentile), medium (25th–75th percentile) and high (upper 75th percentile; **Fig. 3g, Fig. S3c**, Methods), and performed differential expression analysis between the expression profiles of cells with matched sgRNAs in low-vs. high-diversity regions (Methods). The expression of multiple genes robustly depended on the clonal diversity of the micro-environment across clones, sections and tumors (**Fig. 3h, Fig. S3d**). Hallmark epithelial-to-mesenchymal transition (EMT) markers^52^, such as PDGFRB, COL5A2, TNC, were higher in clones in low entropy regions, while the pleiotropic cytokine GDF15 was found to be particularly upregulated in regions of high clonal diversity. Taken together, PerturbView provides an integral platform for merging single-cell spatial transcriptomics with sgRNA readouts, paving the way for spatial perturbation experiments.

## DISCUSSION

Optical Pooled Screening (OPS) is a versatile tool for dissecting biological processes. However, its practical application has mostly been limited to low-plex phenotypes in cancer cell lines, due to challenges in efficiently and accurately detecting perturbation barcodes. Here, we have addressed these limitations with PerturbView, a novel OPS approach that leverages IVT^25^ to amplify sgRNA barcodes, facilitating their detection across diverse cell types and tissue and their combination with different multiplexed phenotyping assays for RNA and proteins. In addition, PerturbView signal is localized to the nucleus, making it easier to assign perturbations to challenging-to-segment cell types (e.g., neurons, fibroblasts) than conventional OPS.

Our PerturbView vector was adapted from CROP-seq^28^ by replacing the U6 promoter with an engineered chimeric U6/T7 promoter, which enables IVT, while preserving CRISPR efficiency. PerturbView cell libraries are also compatible with Perturb-Seq^3^, enabling matched optical and molecular profiling.

Our PerturbView screen of the NFκB pathway in BMDM cells is, to the best of our knowledge, the first OPS in any primary cell. Because PerturbView has high detection efficiency, it significantly reduces the number of cells required for screening, a major consideration with primary cells. Moreover, compared to an earlier OPS of the NFκB pathway in a cancer cell line^13^, our screen in primary immune cells shows key context-dependent biological differences, underscoring the importance of screening in a relevant biological context.

PerturbView facilitates multiplexed and multi-modal OPS, as well as *in vivo* screens in animal models. While RNA and protein phenotyping before *in situ* sequencing compromised conventional OPS performance, in PerturbView, we successfully executed IVT post-phenotyping, thereby decoupling RNA detection from potential degradation and diffusion before OPS barcoding. In principle, any non-destructive *in situ* phenotyping method can be used prior to PerturbView, including highly-multiplexed techniques for protein detection, DNA and RNA FISH, and epigenetic modulators. Similarly, PerturbView can be used for pooled tissue screens of CRISPR perturbations with spatial-omic readouts, which were previously hampered by inefficient RNA barcode detection (with ISS).

FISH-based detection of sgRNAs and barcodes has recently emerged as an alternative to ISS for reading out multiplex screens^12,32,54^. These methods generally either directly detect an sgRNA with a FISH probe or detect an expanded FISH barcode often proximal to the sgRNA of interest. FISH methods excel at the rapid identification of barcode sequences but have several limitations: expanded FISH barcode proxies for sgRNA risk recombination during lentiviral production (resulting in high misidentification rates)^55,56^, and necessitate more complex cloning procedures; and directly probing against an sgRNA necessitates one FISH probe per sgRNA, severely limiting the multiplex capacity and potentially risking high crosstalk between FISH probes. In any case, we anticipate that integrating the PerturbView construct design with FISH-based readouts will substantially improve the fidelity and speed of future FISH-based CRISPR screens.

Future developments to PerturbView can further increase the method ‘s detection efficiency and accuracy, expand compatible phenotypic assays, and improve the throughput of perturbation experiments in tissues. While barcode diffusion results in minimal tradeoffs in sensitivity and specificity, improvements can be made to better anchor IVT-generated RNA to reduce diffusion. Tissue perturbation experiments may require large numbers of cells to decipher the effects of perturbations in a complex tissue microenvironment. Future versions of PerturbView can substantially increase their throughput through optimizations to work in 3D by utilizing tissue clearing^57^ and deep fluorescence imaging technologies^58^.

In conclusion, PerturbView provides a key tool for optical pooled screening, with unprecedented sensitivity, accuracy and breadth, across cell lines, primary cells and *in vivo* in animal models. PerturbView ‘s compatibility with multiplexed phenotyping and spatial-omic readouts will enable comprehensive dissections of the impact of genetic perturbations and tracking of cell states across health and disease.

## METHODS

### Cell culture

HEK 293T, 3T3, L929, Raw 264.7, DLD-1 and A549 cells were all obtained from ATCC and maintained in DMEM supplemented with 10% Fetal Bovine Serum (FBS; v/v), 100 units/ ml penicillin, 100 μg/ml streptomycin and 1X GlutaMAX.

IMR-90 fibroblasts were obtained from ATCC (ATCC CCL-186) and maintained in EMEM (ATCC 30-2003) supplemented with 10% (v/v) FBS.

For iPSC-derived neurons, iPSCs (ALSTEM, iP11N) were differentiated using a strategy combining NGN2 programming and small molecule patterning optimized from a previously published protocol^59^. On day 7 of differentiation, cells were lifted from the flask and plated at 40,000 cells per well in poly-d-lysine and iMatrix-coated 96-well plates. On day 8, cells were transduced by adding appropriate volumes of lentiviral supernatant overnight to achieve an MOI of < 0.3. Cells were then subjected to antibiotic selection by puromycin at 1 μg/ml on day 11 for 3 days. At day 18, cells were fixed for 30 minutes in 4% paraformaldehyde in 1X PBS for *in situ* sequencing.

iAstrocytes were generated from iPSCs (ALSTEM, iP11) based on a published protocol^60^. Immature astrocytes harvested from astrospheres were plated at 10,000 cells per well in matrigel coated 96-well plates for further maturation. A day later, cells were transduced by adding appropriate volumes of lentiviral supernatant overnight to achieve an MOI of < 0.3. Cells were selected by puromycin treatment at 2 μg/ml at day 5 of maturation for 3 days and fixed at day 8 with 4% paraformaldehyde in 1X PBS for 30 minutes.

Immortalized macrophages (ER-Hoxb8-immortalized murine myeloid progenitor cells^61,62^) were established from Cas9 transgenic mice^63^. Cells were cultured in RPMI 1640 medium supplemented with 10% (v/v) FBS, 20 ng/mL murine granulocyte-macrophage colony-stimulating factor (GM-CSF, eBioscience), and 1 mM β-estradiol (Millipore Sigma). Cells were transduced by spinfection at 3200 × g at 32°C for 45 min with 5 μg/ml polybrene, cultured for another 2 days and then selected with 10 μg/ml puromycin. Cells were differentiated into macrophages in DMEM supplemented with 10% (v/v) FBS, and 20% (v/v) L929-conditioned medium at 37°C with 5% CO_2_ and were re-plated to 96-well plates at 20,000 cells per well on day 5 for experiments.

Bone marrow was harvested from tibias and femurs of 12–18 weeks old Cas9-eGFP transgenic mice^64^. After red blood cells lysis (ACK lysis buffer, GIBCO A1049201), 20 M cells were plated in BMDM media [DMEM high glucose, 10% Tet-Negative heat-inactivated FBS, 1X GlutaMAX, 100 U/ml penicillin-streptomycin and 100 ng/ml recombinant murine M-CSF (Genentech media facility)] at a density of 0.8 M cells/ml in 150-mm non-TC treated dishes (Corning). Cells were transduced on day 3 by adding a single lentivirus or lentiviral pool supernatants overnight in fresh BMDM media. Transduced cells were selected with 5 μg/ml puromycin from day 8 to day 11.

T cells were isolated using the pan-T cell isolation kit (Miltenyi) and frozen in FBS supplemented with 10% DMSO. Following thawing, T cells were resuspended in T cell media (X VIVO 15 + 5% FBS + 500 U/mL IL-2) to a concentration of 1×10^6^ cells/mL, and stimulated with anti-human CD3/CD28 CTS Dynabeads (ThermoFisher Scientific, catalog no. 40203D) at a 1:1 cell:bead ratio. Approximately 18 hours post-stimulation, cells were transduced with Lenti-Cas9 (2.5% v/v); 42 hours after stimulation, cells were transferred to a 96-well plate and transduced with 0.25% v/v of CD81 sgRNA virus. If sgRNA constructs contained a puromycin selection cassette, cells were then selected with 2 μg/mL puromycin on day 3. Over the next few days, cells were expanded and maintained by adding media or splitting as necessary. On day 6, CRISPR-mediated CD81 knockout was assessed by flow cytometry. Cells were then plated onto CD3 antibody coated lysine surfaces, allowed to adhere for 15 minutes at 37°C before 4% PFA fixation followed by ISS.

All cells were cultured at 37°C and 5% CO_2_ in a humidified incubator.

### Animal experiments

All protocols involving animals were approved by Genentech ‘s Institutional Animal Care and Use Committee, in accordance with guidelines that adhere to and exceed state and national ethical regulations for animal care and use in research. All mice were maintained in a pathogen-free animal facility under standard animal room conditions (temperature 21 ± 1°C; humidity 55–60%; 12 h light/dark cycle).

### Imaging

Imaging was performed on a Nikon Ti2 microscope equipped with Nikon lenses, a Hammamatsu Fusion BT camera and a Lumencor CELESTA light engine.

*In situ* sequencing cycles were imaged using either a 10X 0.45 NA CFI Plan Apo Lambda or a 4X 0.20 NA CFI Plan Apo Lambda objective with the CELESTA-DA/FI/TR/Cy5/Cy7 (Semrock) excitation filter and FF01-391/477/549/639/741 dichroic unless noted otherwise. Laser excitation wavelength and emission filter are set up for each channel: DAPI (408 nm, FF01-441/511/593/684/817); base G (545 nm, FF01-575/19); base T (545 nm, FF01-615/24); base A (637 nm, FF01-680/42); base C (637 nm, FF01-732/68).

For p65 phenotyping, the following conditions were used for imaging IF signal with a 10X objective: DAPI (365 nm, FF01-433/24) and CF430 (446 nm, 5%, ex: FF01-432/523/702, em: FF01-470/28, dm: FF459/526/596-Di01).

HCR FISH phenotypic images were acquired with a 20X 0.75NA CFI Plan Apo Lambda objective with the following conditions: DAPI (365 nm, FF01-441/511/593/684/817); AF594 (561 nm, FF01-615/24); AF647 (637 nm, FF01-680/42), AF750 (748 nm, FF01-792/64).

IBEX Immunofluorescence images were acquired with a 20X 0.75NA CFI Plan Apo Lambda objective the following conditions: DAPI (365 nm, FF01-441/511/593/684/817); CF430 (446 nm, ex: 432/523/702, em: FF01-470/28, dm: 459/526/596); GFP (477 nm, FF01-511/20); AF532 (545 nm, ex: FF01-432/523/702, em: FF01-563/9, dm: 555-Di03); AF647 (637 nm, FF01-680/42); AF750 (748 nm, em: FF01-792/64).

For imaging time comparison between the conventional protocol with 10X magnification and PerturbView with 4X magnification, the following exposure was used: DAPI (200 ms), base G (200 ms), base C (200 ms), base A (200 ms), base T (800 ms). 10% and 4% overlapping regions were acquired for 10X and 4X objective lenses, respectively, to use the same physical distance for the image registration. The same exposure was also used for the frameshift reporter assay at 4X magnification.

### Lentiviral transduction

HEK293T cells were seeded into 6-well plates at a density of 1 million cells per well. The following day, cells were transfected with the expression plasmid, delta8.9 and pCMV-VSV-G at a molar ratio of (1:2.3:0.2) using Lipofectamine 3000 (ThermoFisher Scientific L3000015). Opti-MEM (Gibco 31985062) was exchanged to culture media after 4 hours. Viral supernatant was harvested 2 days after transfection and filtered through a 0.45 μm filter. For infection of BMDMs, viral supernatant was further concentrated by adding Lenti-X concentrator (Takara 31231) and centrifuging at 4°C for 1 hour. LentivirusL titer was quantified before the screens using CellTiter-Glo kit (Promega G7570).

### *In situ* sequencing

Cells were fixed with 4% paraformaldehyde (Electron Microscopy Sciences 15714) in PBS for 30 minutes, and washed three times with PBS with 0.05% Tween (PBST). Cells were permeabilized with 70% ethanol for 30 minutes, followed by 75% volume exchange with PBST three times to prevent drying and two complete PBST washes. Afterward, heat-decrosslinking was carried out in 0.1 M sodium bicarbonate and 0.3 M NaCl in water at 65°C for 4 hours (cell culture) to 24 hours (tissue). It is important to use glass-bottom plates that tolerate high heat (Cellviz, P96-1.5H-N or P12-1.5H-N). After three washes with PBST, IVT was conducted using the HiScribe® T7 Quick High Yield RNA Synthesis Kit (NEB E2050S) - 1x T7 RNA Polymerase Mix, NTP Buffer Mix at 10 mM with the addition of RiboLock RNase inhibitor (ThermoFisher Scientific EO0384) at 0.4 U/μl at 37 °C for 12 hours. For control samples without IVT, fixed cells were kept in PBST with 0.4 U/μl Ribolock at 4°C for less than 48 hours after permeabilization.

A primer crosslinking protocol was adopted from Labitigan et al.^21^ for all data, except **Figure 1b-c** which uses the original OPS protocol^13^. An amine-modified primer was hybridized at 1 μM in PBST for 30 minutes at room temperature. After the hybridization and before RT, cells were fixed with 3.2% paraformaldehyde and 0.1% glutaraldehyde (Electron Microscopy Sciences 16120) in PBST for 30 minutes. The reaction was quenched by adding 0.2 M Tris-HCl (pH 8), and cells were washed three times with PBST. The reverse transcription reaction mix (1x RevertAid RT buffer, 250 μM dNTPs (NEB N0447L), 0.2 mg/mL recombinant albumin (NEB B9200S), 1 μM RT primer, 0.4 U/μL Ribolock RNase inhibitor, and 4.8 U/μL RevertAid H minus reverse transcriptase (ThermoFisher Scientific EP0452) was added into samples, followed by incubation at 37°C overnight.

After reverse transcription, cells were washed three times with PBST and fixed again in 3.2% paraformaldehyde and 0.1% glutaraldehyde in PBST for 30 minutes. Cells were then washed with PBST six times to remove residual dNTP. After this step, cells were incubated in gap-fill reaction mix (1x Ampligase buffer, 0.4 U/μL RNase H (Enzymatics Y9220L), 0.2 mg/ mL recombinant albumin, 100 nM padlock probe, 0.02 U/μL TaqIT polymerase (Enzymatics P7620L), 0.5 U/μL Ampligase (Lucigen A3210K) and 50 nM dNTPs) for 5 minutes at 37°C and 90 minutes at 45°C. Cells were washed with PBST three times, and rolling circle amplification was performed by adding the reaction mix (1x Phi29 buffer, 250 μM dNTPs, 0.2 mg/ mL recombinant albumin, 5% glycerol, and 1 U/μL Phi29 DNA polymerase (ThermoFisher Scientific EP0091) and incubating at 30°C for >20 hours.

*In situ* sequencing steps were conducted according to the OPS protocol^13,30^ with the primer sequences listed in **Table S2**.

### Assessment of editing efficiency

For determining editing efficiency, a targeting sgRNA was transduced by lentivirus to Cas9-expressing cells. Next, cells were cultured for 4 days and then fixed in 4% paraformaldehyde in PBS for 30 minutes. Cells were washed twice with PBS and then incubated with fluorophore-conjugated antibodies (CD81 antibody BioLegend 349501 or Cd44 antibody Biolegend 103023) for 1 hour followed by a PBS wash. Antibody signal was measured by flow cytometry (Sony, MA900 Multi-Application Cell Sorter) gated on cells based on forward and side scatter.

### PerturbView signal quantification

Cell models were fixed in 4% paraformaldehyde. After 30 minutes, cells were washed with PBST three times. Plates without PerturbView were reserved at 4°C in PBS + 0.4 U/μl Ribolock while decrosslinking and IVT protocols were carried out. Ethanol permeabilization and reverse transcription were performed within 72-hour post-fixation.

Because the massive signal amplification of PerturbView poses a challenge in fitting read intensities within the optics’ dynamic range when using the same exposure setting with the standard OPS protocol, a 4-fold longer duration of exposure was used for non-PerturbView images of iAstrocytes, fibroblasts and iNeurons.

### Frameshift reporter assay

Cas9-expressing A549 cells were first transduced with the frameshift reporter^13^ at high MOI by repeating the transduction processes and selected with 300 μg/ml hygromycin (ThermoFisher Scientific 10687010) for 1 week. Five targeting and five non-targeting control sgRNAs were individually cloned into a PerturbView vector, pooled equimolarly, packaged as lentivirus, and transduced into Cas9-expressing A549 cells at low MOI, followed by puromycin selection. Cells were plated 12,000 cells per well in a 96-well plate (CellViz, P96-1.5H-N) and fixed with 4% paraformaldehyde for 30 minutes on the following day. Epitope staining was conducted as previously described^13^, followed by *in situ* sequencing. As described above, 4-fold longer durations of exposure were used for standard OPS than for PerturbView, due to the different dynamic ranges of the two methods.

### Library construction for BMDM screen

For the BMDM library, four sgRNAs were designed per gene for 23 NFκB-related genes^13^, 130 metabolic and signaling genes (**Table S1**) and 10 olfactory receptor genes (as non-expressed gene controls), along with 60 different non-targeting sgRNAs. Spacer sequences were generated by crisprVerse^65^ to design barcodes that can be uniquely identified by 6 and 8 cycles of *in situ* sequencing for NTC and BMDM libraries, respectively. The oligo pool was synthesized by Twist Bioscience, amplified by dial-out PCR^66^ using Ex Premier DNA Polymerase (Takara Bio), cloned into a PerturbView vector with golden gate cloning using BsmBI, transformed into Stellar electrocompetent cells (Takara Bio), and packaged into a lentivirus library while maintaining >1000x coverage at every step.

To validate sgRNA transduction, gDNA was extracted and amplified according to an established protocol^30^. Indexed amplicons were sequenced on an Illumina MiSeq with 150 cycles and the v3 reagent kit (Illumina, MS-102-3001). The reads were aligned to the expected amplicon sequences (N20 corresponding to the spacer region flanked by backbone sequences) using the Burrows-Wheeler alignment tool (bwa-mem)^67^ to produce reads counts for each designed guide.

### NFκB translocation screens

BMDMs were isolated from mice expressing Cas9-EGFP and cultured in DMEM high glucose, 10% FBS, 2 mM glutamine, 1% Pen/Strep, and 100 ng/ml CSF-1. After 2 days, these cells were split into new plates and infected with the guide-bearing lentiviruses at MOI of <0.2. Media was replaced the following day and selection of the infected BMDMs began with 5 μg/ml puromycin. Three days after the selection and media change, cells were plated in a 12-well plate (Cellviz P12-1.5H-N) at a density of 276,000 cells per well. After 24 hours, cells were stimulated with 10 ng/ml TNFα, 2 ng/ml IL1β, 100 ng/ml LPS and H_2_O for 40 minutes before fixation. Cells were fixed in 4% paraformaldehyde in PBS for 30 minutes. Following three PBST washes, cells were permeabilized with 0.2% Triton-X in PBS for 10 minutes. Cells were then blocked with 3% BSA in PBS for 45 minutes, incubated with p65 antibody in PBS at 1:1000 (Cell Signaling Technologies #8242) for 1 hour and washed three times with PBST. Media was replaced with Donkey Anti-Rabbit CF430 IgG (Biotium 20460) in 3% BSA in PBS and incubated for 30 minutes, followed by three washes with PBST. Prior to imaging, media was replaced with 200 ng/ ml DAPI in 2xSSC (saline-sodium citrate) buffer for 10 minutes. After antibody imaging, ISS was carried out beginning from the ethanol permeabilization step.

### Multimodal screens

BMDMs were transduced, selected and plated in the same manner as above. Cells were stimulated with 10 ng/ml TNFα, 2 ng/ ml IL1β for 24 hours. Cells were fixed in 4% paraformaldehyde in PBS for 30 minutes, washed three times with PBST and then permeabilized with 0.2% Triton-X for 10 minutes, followed by three washes with PBST.

To perform HCR FISH, cells were incubated at 37°C for 30 minutes with probe hybridization buffer (30% formamide, 5x SSC, 0.1% Tween), then incubated with primary HCR FISH probes (Molecular Instruments) against Gbp2 (B2, **Table S2**), Ccl4 (B3, **Table S2**) and Ripk2 (B5, Molecular Instruments) diluted at 1:250 ratio in probe hybridization buffer with 0.4 U/μL Ribolock at 37°C for 4 hours. Samples were then washed with HCR probe hybridization buffer four times, each for 5 minutes at 37°C. Next, samples were incubated in HCR amplification buffer (5x SSC, 0.1% Tween) at room temperature for 30 minutes. HCR amplifiers were separately heated at 95°C for 90 seconds and then cooled to room temperature in the dark for 30 minutes. HCR amplifier mix (Molecular Instruments, B1-488 h1/h2, B2-594 h1/ h2, B3-647 h1/h2 diluted at 1:125 in amplification buffer with 0.4 U/μL Ribolock) were added to the sample, followed by incubation at room temperature for 2 hours. Excess amplifiers were removed by washing five times with HCR probe amplification buffer. An imaging buffer (200 ng/mL DAPI in 2X SSC with) with 0.4 U/μL Ribolock was then added to the sample, followed by a 10 minutes incubation. After imaging, HCR FISH probes were stripped off by adding HCR stripping solution (80% formamide in 2X SSC) and incubated at 37°C for 30 minutes, followed by three washes at 37°C with HCR stripping buffer, each for 5 minutes, followed by three PBST washes.

To perform immunofluorescence, cells were blocked with 3% BSA in PBS for 45 minutes and then incubated with fluorophore direct-conjugated antibodies: Irf-1 conjugated with CF430 (Biotium #92118) with maleimide chemistry (Cell Signaling Technology #8478), Nos2 Alexa Fluor 532 conjugate (ThermoFisher Scientific, 58-5920-80), p65 Alexa Fluor 647 conjugate (Cell Signaling Technology #8801) and phospho-rpS6 Alexa Fluor 750 conjugate (Cell Signaling Technology #62788) in 3% BSA in PBS at 1:75, 1:200, 1:200, 1:200 ratio respectively for 1 hour. After three washes with PBST, samples were placed in 200 ng/ ml DAPI in 2xSSC (saline-sodium citrate) buffer and incubated for 10 minutes before imaging. After imaging, the fluorescent signal was bleached by a freshly made 1 mg/mL lithium borohydride in H_2_O for 30 minutes, followed by three PBST washes^27^. Afterward, *in situ* sequencing was carried out from the ethanol permeabilization.

### Tissue experiments

DLD-1 cells (ATCC CCL-221) were transfected with the NTC library with polybrene. Transfected cells were selected and maintained with 2 μg/ml puromycin. 1.5 million transfected cells were subcutaneously implanted into each 8–12 weeks old female NCR nude mouse (Taconic).

Mice were anesthetized for manual restraint with isoflurane gas to facilitate injection and minimize stress to the animal. As soon as mice were immobile, they were removed from the chamber and placed on a nose cone delivering the anesthetic gas. The injection site area was cleaned with alcohol. Using forceps, the skin over the inoculation site was lifted and a sterile 25 G needle attached to a 1.0 ml tuberculin syringe was inserted through the skin into the subcutaneous space. An inoculum dose of at most 200 μl was deposited. The needle was then held in place for approximately 3 seconds before being carefully withdrawn. The mouse was returned to its cage and monitored until fully recovered from anesthesia.

Two weeks after implantation, DLD-1 cells developed into subcutaneous tumors of ∼250 mm^3^. Individual xenografts were harvested and halved, with half the tumor chunk embedded in OCT and frozen on dry ice immediately and the other half fixed in 10% neutral buffered formalin for 20 hours at 4°C, processed and embedded in paraffin. Fresh frozen and FFPE tumor blocks were cut into 10-μm and 5-μm thick tissue sections, respectively, placed on Superfrost Plus Microscope glass slides or Xenium slides, and stored at –80°C for fresh frozen tissue and 4°C for FFPE tissue.

Fresh frozen samples were taken from –80°C and fixed with 4% paraformaldehyde for 30 minutes in a 50 ml falcon tube. Slides were washed three times by transfer to new 50 mL tubes filled with PBS for 5 minutes each. Each slide was transferred to a tube filled with 200 ng/ml DAPI in PBS and mounted on a SecureSeal hybridization chamber to acquire nuclear images. After imaging, slides were transferred to a tube filled with decrosslinking buffer (0.1 M sodium bicarbonate and 0.3 M NaCl in water) and incubated at 65°C overnight. After PBS washes, slides were mounted on Xenium cassettes (10X Genomics, PN-1000566). Downstream reactions, including IVT, RT, gap-filling and RCA, took place in the Xenium cassettes until the *in situ* sequencing step. Then, a SecureSeal Hybridization chamber was mounted for *in situ* sequencing and imaging. *Z*-stack images were acquired –8 to 20 μm from the tissue plane with 4 μm steps.

FFPE sections were first deparaffinized and de-crosslinked according to the Xenium FFPE protocol^51^. Briefly, FFPE sections were incubated at 60°C for 2 hours, equilibrated to room temperature for 7 minutes, and immersed in xylene and incubated for 10 minutes twice. FFPE sections were then immersed in an ethanol series of 100%, 100%, 96%, 96% and 70% for 3 minutes for each incubation. Finally, sections were rinsed in nuclease-free water for 20 seconds and assembled into Xenium cassettes. Each section was de-crosslinked with 300 μl of Xenium decrosslinking buffer and incubated at 80°C for 30 minutes, 22°C for 10 minutes, and washed with PBST for 1 minute twice before acquiring DAPI images. PerturbView IVT and ISS protocols were performed as previously written.

### Preprocessing of cell screens

For p65 screens, flatfield references were collected experimentally by diluting the CF430 antibody by 1000-fold in PBS, and 49 images were captured at different locations in a 12-well plate. Background references, including autofluorescence from media, were also collected by imaging PBS in the same manner. The median was computed for each pixel to represent flatfield image Ƒ and background image Ɓ. For each image İ, the correction was performed by (İ-Ɓ/Ƒ) μƑ, where μƑ is a mean of Ƒ and works as a scaler. The negative values were clipped to 0.

For multimodal screens, 500 images were randomly sampled, and the flatfield and darkfield images were computed using BaSiC68 for background correction.

### Image registration in cell screens

For multimodal screens, phenotypic images were acquired using 20X magnification for resolution and *in situ* sequencing was acquired at 10X magnification for speed. Due to a jitter introduced between imaging cycles and the chromatic aberrations between objective lenses, a series of alignments was performed. First, for each well and cycle, images were stitched together using ASHLAR69 to represent a whole-well image. Composite well images were then resized 2X by nearest-neighbor interpolation to match the pixel size of phenotyping images. Images across cycles were then cropped to the same size (minimum image size obtained from each well across cycles). These whole-well images across cycles were then coarsely aligned by ASHLAR and split into smaller tiles of 5000×5000 pixels. Affine transformation followed by optical flow-based registration was performed using microaligner^70^ for further linear and non-linear alignment.

### Images segmentation in cell culture

To extract relevant information from cells and nuclei, image segmentation was performed in two sequential steps. First, nuclei were segmented using Stardist^71,72^, and the resulting nuclei served as seeds for cytoplasmic segmentation using watershed. Segmentation information was stored in Zarr format for use along the pipeline.

### Spot detection in cell culture

An *in situ* sequencing analysis pipeline was adopted and modified from a prior study^18^. Spot detection was executed to identify the locations of *in situ* sequencing foci. Aligned images were log-transformed using a Laplacian-of-Gaussian filter with a sigma of 1. All other parameters of the scipy function ‘ndi. gaussian_laplace’ were left as default. The resulting filtered images were then subjected to a max filter with a neighborhood size of 3 pixels. Subsequently, the standard deviation over cycles was calculated for the mean of all channels. The latter array was used to identify local maximum within a neighborhood of 5 pixels. These maxima represent the location of ISS spots that are later decoded during base calling. Spot intensities were extracted and corrected for crosstalk by linear transformation as described previously^13^.

### Base calling and barcode identification

Reads were identified by calling the base with the highest intensity at each spot identified in the first step, utilizing an adhoc quality score per sequencing call, based on the highest and second-highest intensity channels, calculated as:

A table containing barcode information and associated quality scores was then used to assign the most common barcode reads to each cell. The resulting table includes all cells with sequencing reads, listing the top two most common barcode sequences for each cell.

### Phenotypic feature extraction

A table of cellular and nuclear phenotypes was generated based on the previously segmented phenotypic images. Intensity features such as percentiles (0.25, 0.75), median absolute deviation (MAD), maximum, median and mean intensity, intensity standard deviation, and total intensity (sum), were calculated using the region_props module of scikit-image package v0.20.073. Other features included peak intensity and locations, number of peaks, intensity distribution (mean intensity and radial coefficient of variation for each concentric bin), edge intensity (min, mean, max, sum, std along edge pixels), weighted center of mass and mass displacement, and the Hu moments (translation, scale and rotation invariant) of the intensity image.

### Merging of ISS and imaging phenotypes

Results from *in situ* sequencing were merged with phenotypic imaging data based on their common cell label. The resulting table contained all the features listed above, for each channel of the phenotype image, along with the label of each cell, well identity, barcodes identified, and barcode associated information (ID, gene, chromosome, guide name, etc.).

### Post-processing of screening data

As multimodal data required complex image registration steps and potentially contains alignment errors, the alignment was evaluated by checking the overlap between the segmentation mask and DAPI channel image from each cycle. A rectangular bounding box was defined for each cell based on the segmented nuclei. The pixel-wise correlation inside the bounding box between the first cycle and each later cycle was calculated. The correlation is close to 1 if the DAPI image and nuclear mask overlaps, indicating little to no alignment error. Cells were removed from analysis if the minimum correlation coefficient across cycles were less than 0.9. Cells were further filtered out by removing small nuclei (area < 400 pixels).

### Channel threshold calculation for estimation of positive and negative populations

To address the different background signal in each condition and cell line, a method was developed to compute distinct thresholds, by adopting an approach used in HTODemux^74^, where barcode assignment also requires a background estimate for each batch and cell hashing oligo. First, a local maximum was computed within the corresponding cell labels (cell segmentation for standard OPS and nucleus segmentation for PerturbView) and those pixels were collected as potential reads (i.e., each pixel position contains a vector of length 4 corresponding to each channel). For efficiency, the number of pixels for the following computation was limited to 25,000. Then, the data were centered-log-ratio normalized and clustered with K-medoids clustering with K=C+1, where C=4, representing four bases and background pixels for a mixed library, or C=1 representing one base and background pixels for a single construct. Next, for each channel individually, and for each cluster, the average channel intensity was computed and the cell population that belongs to the bottom three clusters (as two of the channels have high crosstalk for C=4) or one cluster (C=1) was selected to represent the background population. The top 0.5 percentile of cells were excluded as outliers, and a negative bimodal distribution was fit to the obtained cluster. The 99th percentile of the fitted distribution was taken as the threshold for each channel. Each channel was thresholded accordingly and the remaining spots were segmented as reads.

### Frameshift reporter assay analysis

The existing algorithm for base calls in OPS^13,18^ relies on the assumption of the relative scale of each channel and often does not work well for the single cycle *in situ* sequencing images. The new HTODemux-based thresholding algorithm described above was therefore used for spot detection instead. Pixels were then corrected for crosstalk by linear transformation^13^ and normalized by *Z*-score transformation for each channel. The base was determined based on the maximum intensity observed across all channels.

To classify each cell into HA-positive and HA-negative, a two-component Gaussian mixture model was fitted to the distribution of mean HA intensity. A probability threshold of 0.95 was used to remove uncertain cells from data. Note that false positives (HA-negative cells with a targeting guide) could be attributed not only to barcode diffusion but other causes, such as reporter variability, multiple integrants and recombination^56^ or downstream analysis errors in segmentation and base calling.

Sensitivity and precision were computed in a standard manner: sensitivity = TP/(TP+FN); precision = TP/(TP+FP)

### p65 screen analysis

For each well, single-cell p65 intensities were normalized using the robust *Z*-score transformation based on the cell population carrying non-targeting control sgRNA, and aggregated over replicate wells. One, two and three-well replicates were collected for LPS, TNF and IL1β stimulations, respectively. For *p*-value calculations, following prior work^13,18^, for each guide, 100,000 null distributions were obtained by randomly sampling the same number of median intensity values from the non-targeting control population, and then a gene-level null distribution was obtained by sampling 100,000 times from the corresponding guide-level null distributions. This empirical cumulative null distribution was then compared to the median p65 intensities for each gene to determine *p* values. To handle non-targeting controls as a ‘dummy’ gene, a set of four non-targeting control sgRNAs were randomly grouped together. Benjamini–Hochberg FDR *q*-values were calculated and FDR < 0.05 (two-sided) was used to call hits.

### Multimodal screen analysis

Phenotypic features were extracted from two rounds of imaging as described above. DAPI replicates were excluded from later phenotyping rounds and Cas9-GFP signal was excluded from analysis. 131 features that contained > 2% missing values (NaN or Inf) were removed, and cells containing the remaining missing values were also discarded. Data were then normalized by a well-wise robust *Z*-scoring on the NTC population. Features with NTC IQR less than 0.1 were removed, followed by retaining only 356 features whose standard deviation lies within the range of 0.1 to 5.0. Outlier values with a normalized score >5 were discarded by removing any features with >0.1% outliers as well as any remaining cells with outlier values. Finally, guides with <15 remaining cell counts were also discarded. PCA was applied to keep 95% of the variance, resulting in 55 principal components. The filtering retained 205,488 cells from 215,275 cells.

For gene-level embeddings, PCA features were subjected to sphering based on NTC population^75^, aggregated across cells and re-centered on the mean of the negative controls. The centered features were further aggregated by taking the median across guides. Leiden clustering (resolution=1.0)^76^ was performed on these features while excluding NTC guides. Features, including NTC, were projected to PHATE space using cosine distance for a multidimensional scaling^46^.

For guide embedding, PCA features were transformed by robust *Z*-score normalization based on NTC population. Guide features were aggregated by taking the median across cells and projected into a PHATE space. Gene name labels are noted in **Figure S2f** if at least two guides targeting the same gene were among the same *k* = 5 nearest neighbors in the PHATE space.

Significance was scored by an empirical cumulative distribution function based on leave-one-out cosine similarity^77^. For each guide, cosine similarity was computed against the average of the other three guides targeting the same gene. To generate a null distribution, random guides were chosen and bootstrapped 1000 times for null distribution, and a *p*-value was obtained empirically relative to the null distribution. Benjamini–Hochberg FDR *q*-values were calculated and FDR < 0.05 (two-sided) was used to call hits.

For “channel-wise” significance of each perturbation (per feature), input features were restricted to a set exclusive to each imaging channel. Clustering significant perturbations by the distribution of their scores across channels resulted in a heatmap representing how strongly perturbations affect the phenotypic changes captured in each imaging.

### Tissue PerturbView analysis

Background subtraction and illumination correction were performed using BaSiC68 as described above. The *in situ* sequencing images were sampled from *Z* stack images by maximum-intensity projection. To facilitate nuclear segmentation, the best-focused DAPI images were chosen by the following procedure: First, the images from the *Z* stack were divided into 100×100 blocks, then for each block and plane, the power log–log slope was computed as a metric of blurriness^78^. The least blurry plane was sampled from each block to produce a 100×100 matrix representing the coarse focal plane, and a median filter with a kernel size of 3 was further applied to remove outliers. The matrix was then resized to the original image size by linear interpolation and the resulting X, Y pixel was sampled from the *Z*-stacks accordingly.

Nuclear segmentation was performed using Stardist^71,72^ and expanded 5 pixels by dilation to account for registration errors. Each tissue section contains mouse cells surrounding the tumors and a necrotic region that does not produce barcodes. To obtain relevant statistics from the transduced cells, segmentation masks corresponding to non-necrotic tumor tissue were chosen visually using Napari viewer. Each sample replicate contained at least 27,802 cells for analysis.

### Base-calling of tissue PerturbView

To mitigate jitters between cycles, a maximum filter with a kernel size of 9 pixels was applied to 4 channel images of the *in situ* sequencing. Separately, local maxima of the Laplacian of Gaussian images were computed and aggregated across channels for potential peaks. Peaks within a 5-pixel radius were aggregated by sampling the brightest intensity for each channel. Peaks less than an intensity threshold were filtered out. Peaks were then linked over cycles based on location using trackpy^79^ with a search radius of 10 pixels. Peaks that were present in at least 5 cycles were retained for further analysis. Cross-talk of channel intensities was corrected as described above. Bases corresponding to the maximum intensity across channels were assigned as a read sequence.

### Xenium–PerturbView analysis

500 images were randomly sampled, and the flatfield and dark-field images were computed using BaSiC^68^. After illumination correction, images were stitched together using m2stitch^80^, with the detected stage coordinates used as an initial guess for the tile locations. Each stitched image was then subsequently aligned to the maximum-intensity projection of the Xenium DAPI image using WSIReg^81^ using a rigid transformation followed by a nonlinear spline transformation.

For Xenium transcriptome measurements, cells were filtered if <10 genes were detected. For each cell, reads were normalized such that the sum of reads are 10,000 and log-transformed (X+1). For clonal analysis, barcodes present in <20 cells were discarded. Spatial DBSCAN was performed with eps=50 and min_samples=1 to assign barcodes to cells without ISS signals. The Shannon diversity index of sgRNAs was calculated for each cell based on a window size of 200 μm. Top and bottom quartiles were defined as high and low diversity, respectively. sgRNA clones that contain more than 50 cells for each high and low group were retained for analysis. For each tumor, top 100 highly variable genes were selected and differential analysis was performed between individual sgRNA across high and low diversity using a two-sided Wilcoxon test. Genes were aggregated across clones by combining adjusted *p*-values using Fisher ‘s method. Genes were qualified as hits if they had an adjusted *p*-value of less than 0.05 and were expressed in at least 20% of the population.

## Statistics, Reagents or Animal models

All the relevant information is available in Methods.

## Computing environment and data storage

Computational analyses were conducted in the Genentech/ Roche cloud, Google Cloud Platform (GCP) or AWS cloud environment for scalability. Data, including intermediate and final results, were stored in S3 or Google Cloud buckets.

## Acknowledgments

We thank Leslie Gaffney and Anna Hupalowska for their help with figure preparation.

## Inclusion and Ethics statement

### Data availability

Data that are not included in the paper are available upon request.

### Code availability

All code are available upon request.

---

**Fig. S1.**
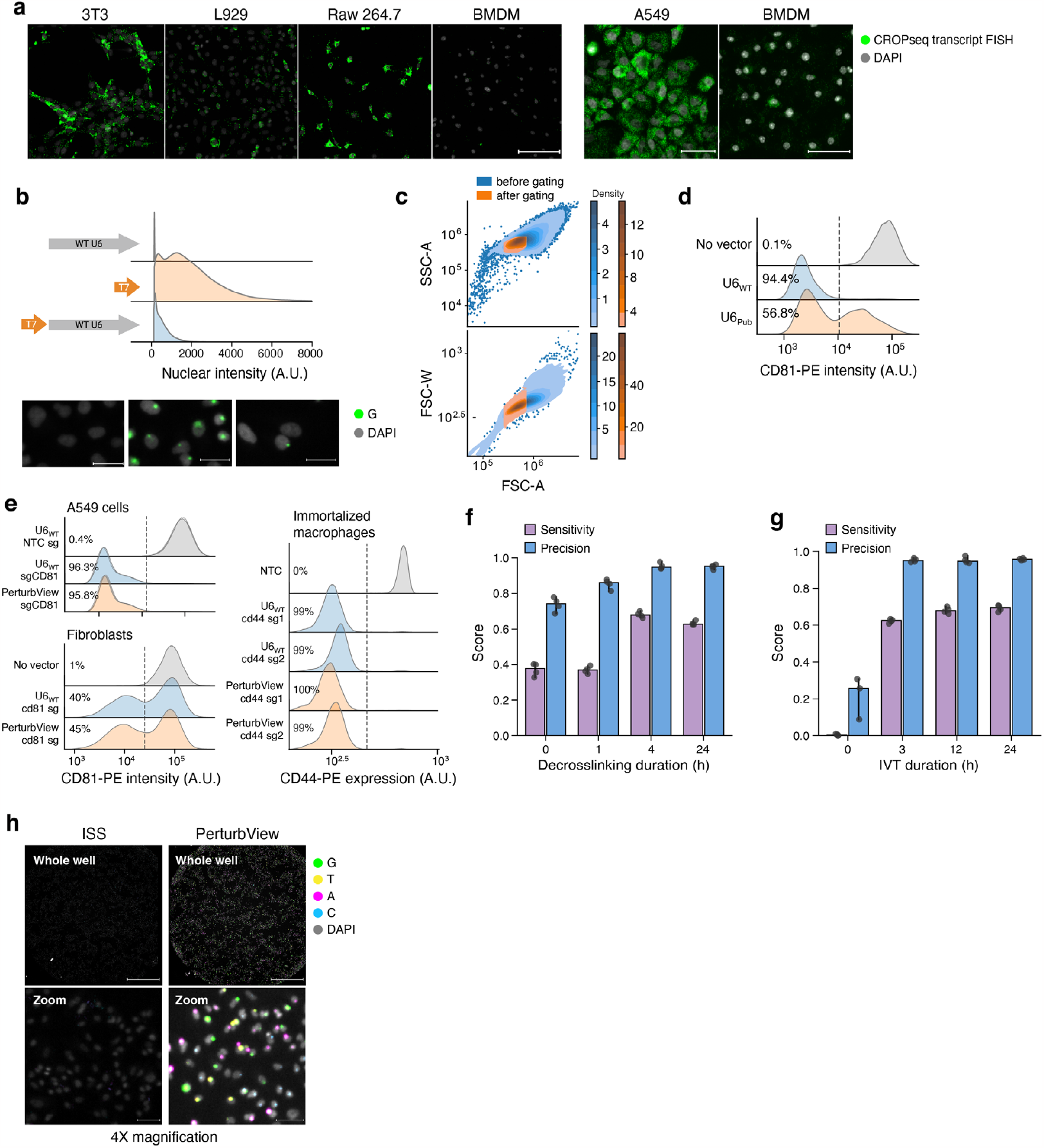
Establishment of PerturbView. (**a**) Varied activity of CROP-Seq vector by HCR FISH in primary cells. Left, HCR FISH of CROP-Seq expressed transcript (green; detected with FISH probes against puromycin and U6 promoter regions; **Table S2**) across different cell lines (scale bar, 200 μm). Right, CROP-Seq transcript density in A549 (left) and mouse BMDM (right) cells (scale bar, 50 μm). All cells were transduced at low MOI and puromycin-selected. (**b**) T7 promoter position dramatically affects IVT-enhanced ISS signal. Distribution of median single cell nuclear intensity following IVT in the original U6 promoter (top; gray, *n* = 62,321 cells pooled from three well replicates), T7 promoter in place of the U6 promoter (orange, *n* = 48,123), and T7 promoter upstream of U6 promoter (blue, *n* = 53,777). Bottom, representative cellular images from each construct (scale bar, 25 μm). (**c**) An example of a gating strategy of flow cytometry data. Top, FSC-A and SSC-A profiles before and after gating. Bottom, FSC-A and FSC-W profiles before and after gating. **(d)** The U6/T7 promoter31 has lower CRISPR editing activity than the unmodified U6 promoter. The distribution of CD81 expression by flow cytometry for each construct. The fraction of knocked-out cells (thresholded by the vertical line) is noted on the left. *k* = 34,950 cells, at least 11,247 cells per condition. **(e)** Editing efficiency of the PerturbView vector is identical to wild-type U6 promoter in A549 cells, immortalized macrophages and fibroblasts. Distribution of target gene expression of each construct in A549 cells (top; CD81, *n* = 3 independent infection replicates combined; 876,505 cells, at least 61,250 cells per replicate and condition), primary IMR90 fibroblasts (bottom; CD81, 17,497 cells, at least 5,524 cells per condition) and immortalized macrophages (right; CD44, sg1 and sg2: separate guide replicates; 126,093 cells, at least 20,844 cells per condition). The fraction of knocked-out cells (thresholded by the vertical line) is noted on the left. (**f,g**) Decrosslinking and IVT duration impact sensitivity and precision of barcode detection. Mean Sensitivity (purple) and precision (blue) by at different durations of decrosslinking (e, *n* = 4 well replicates (dots), at least 9,393 cells per replicate) or IVT (*n* = 4 well replicates (dots), at least 9,387 cells per replicate). Error bars: SD. (**h**) PerturbView enables in situ sequencing at 4X magnification. Left, Whole well (top; scale bar, 1 mm) or zoomed image (bottom; scale bar, 50 μm) representative image acquired at 4X magnification with either conventional ISS (left) or PerturbView (right). All images are contrasted identically. Heat decrosslinking prior to PerturbView results in increased nuclear DAPI intensity.

**Fig. S2.**
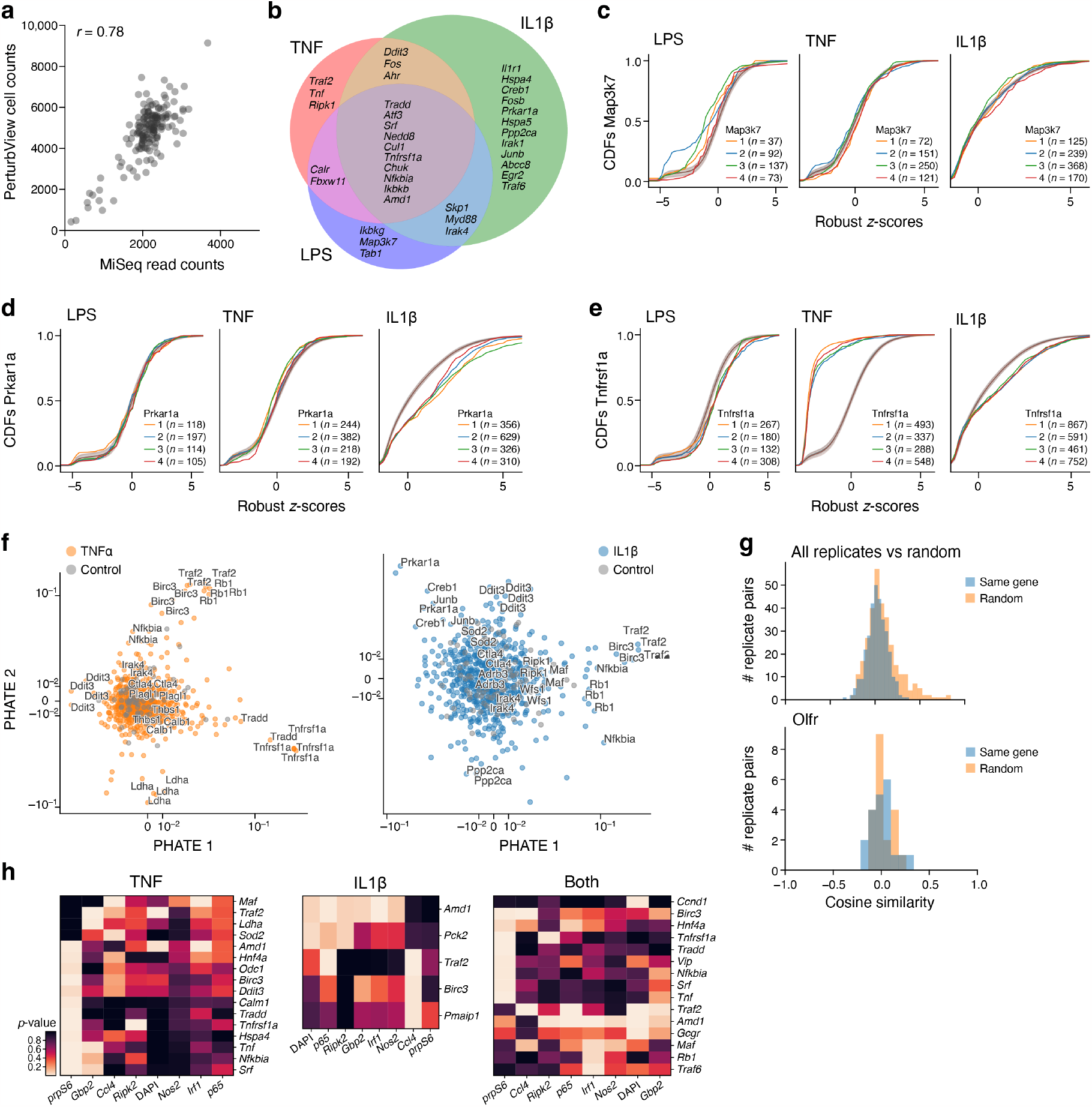
Context and phenotype specific perturbation effects in PerturbView screen in primary immune cells. (**a**) Agreement in sgRNA detection in PerturbView and NGS. PerturbView ISS cell counts (*y* axis) and NGS read counts of the cell library (*x* axis) for each perturbation barcode (dot). Top left: Pearson ‘s r. (**b–e**) Shared and stimulation-specific perturbation effects. (**b**) Intersection of hit genes between TNFα, IL1β, and LPS PerturbView screens (circles) of NFKB translocation in primary BMDMs. (**c–e**) Cumulative distribution functions (CDFs) of robust *z*-scores for p65 nuclear intensity in response to LPS (left), TNFα (middle), or IL1β (right) for each of four guides (colored curves; *n* = cell number per guide) targeting the genes Map3k7 (**c**), Prkar1a (**d**), and Tnfrsf1a (**f**), compared to cells with non-targeting or non-essential controls (Methods; gray line (combined); shading standard deviation of robust *z*-scores). (**f,g**) Agreement in phenotypic profiles of different guides targeting the same gene. f, PHATE embeddings of RNA/protein joint profiles (dots) for each sgRNAs in the TNFα (left) or IL1β (right) screen targeting a gene (color) or controls (grey). Gene names are shown for each guide whose 5 nearest neighbors contain another guide targeting the same gene. (**g**) Distribution of cosine similarity of the phenotypic profiles derived from joint RNA/protein profiling for random pairs of guides (“random”, blue) and either guides targeting the same gene across all genes (orange; “same gene”, top) or targeting Olfr genes (orange “same gene”, bottom). (**h**) Contribution of different molecular phenotypes to perturbation impact. Impact score (*p*-values, color bar) for each perturbed gene (row) in the TNFα (left), IL1β (middle) screen, or both (right), when assessed only based on one imaging feature (RNA, protein or DAPI; columns). Rows and columns are clustered by hierarchical clustering.

**Fig. S3.**
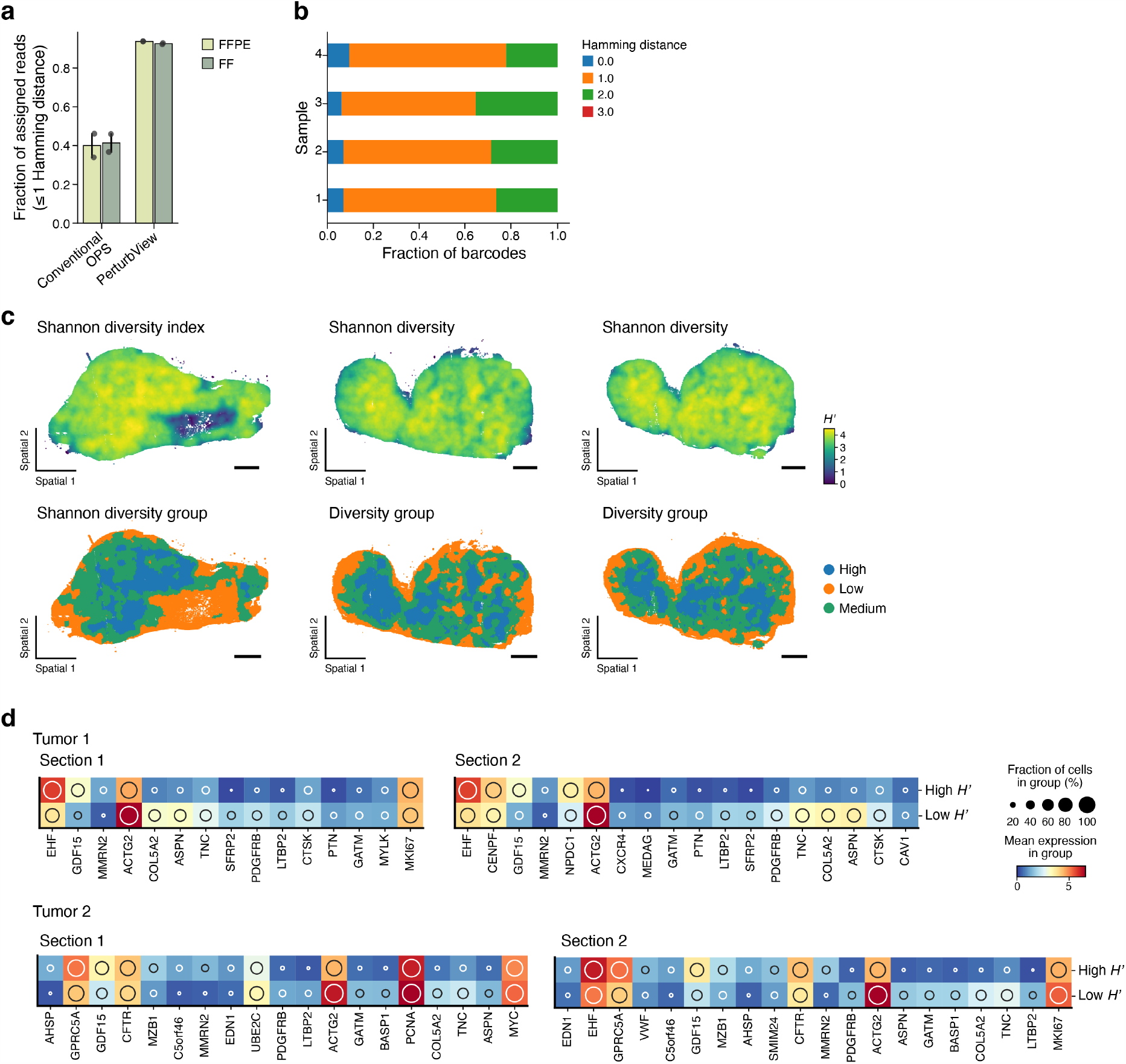
PerturbView in tissue. (**a,b**) High barcoding decoding accuracy in tissue sections with PerturbView. (**a**) Barcode mapping rate (*y* axis, fraction of assigned reads at a Hamming distance ≤ 1 to the pre-defined sgRNA lookup table) for conventional ISS and PerturbView (*x* axis) in FF and FFPE tumor tissue (*n* = 4). (**b**) The Fraction of barcodes (*x* axis) mapped at different Hamming distances (colors) in each of four samples (*y* axis). (**c,d**) Expression changes associated with clonal diversity. (**c**) Tumor sections profiled (scale bar, 1 mm), colored by Shannon diversity (top) or corresponding diversity groups (bottom) for all analyzed sections aside from the one shown in **Figure 3g**. (**d**) Genes (columns) that are differentially expressed between regions with high and low Shannon diversity (rows) for each tumor section (*n* = 2 consecutive sections per tumor).

